# MedSAGE: Bridging Generative AI and Medicinal Chemistry for Structure-Based Design of Small Molecule Drugs

**DOI:** 10.1101/2025.05.10.653107

**Authors:** Alexander S. Powers, Tianyu Lu, Rohan V. Koodli, Minkai Xu, Siyi Gu, Masha Karelina, Ron O. Dror

## Abstract

While generative AI is transforming the *de novo* design of proteins, its effectiveness for structure-based design of small molecules remains limited. Current methods, including diffusion models, often produce small molecules with difficult-to-synthesize structures, poor medicinal chemistry properties, and limited target selectivity. To address these limitations, we introduce MedSAGE, a novel generative AI framework that adapts diffusion models specifically for *de novo* small-molecule design. Rather than using atoms or strings, we develop a novel representation for generating molecules using fragments relevant for medicinal chemistry. Chemical and geometric information characterizing these fragments is embedded in a smooth and interpretable latent space. We also develop an algorithm to optimize the connectivity between generated fragments while preserving chemical validity and synthesizability. In a benchmark of multiple methods across 25 therapeutically relevant protein targets, MedSAGE achieved state-of-the-art performance, producing synthesizable, drug-like molecules with predicted affinity and selectivity closely matching known drugs and drug candidates. Compared to large-scale virtual screening, MedSAGE produced molecules with high predicted affinity over 100 times more efficiently. Our results demonstrate that MedSAGE is already practically useful and paves the way for next-generation tools in structure-guided drug design.

## Introduction

Structural biology is advancing at an extraordinary pace; advances in experimental and computational techniques have enabled the rapid determination of detailed structures for a wide variety of proteins, including many targets of therapeutic interest^1–3^. Despite advances in structure determination, using protein structures to design effective therapeutics remains a complex and costly challenge. Recently, generative artificial intelligence (AI) has emerged as a promising approach for structure-based drug discovery. Diffusion models, a class of generative AI methods^4^, have delivered particularly impressive results recently for the *de novo* design of antibodies and other proteins^5–7^.

However, current generative AI techniques—including diffusion models—have yet to provide robust solutions for structure-based design of small-molecule drugs, the modality encompassing the majority of drugs and new drug approvals. Despite notable theoretical advances^8–12^, these methods continue to fall short in practical utility. Recent *de novo* molecule generation methods, with state-of-the-art performance in the field, still produce molecules that fail to meet basic medicinal chemistry criteria or would not be chemically stable in a physiological environment (Fig. 1b). Reliably generating molecules with critical properties like high affinity^13^, selectivity^14^, and drug-like physicochemical properties^15,16^ remains elusive.

**Figure 1:**
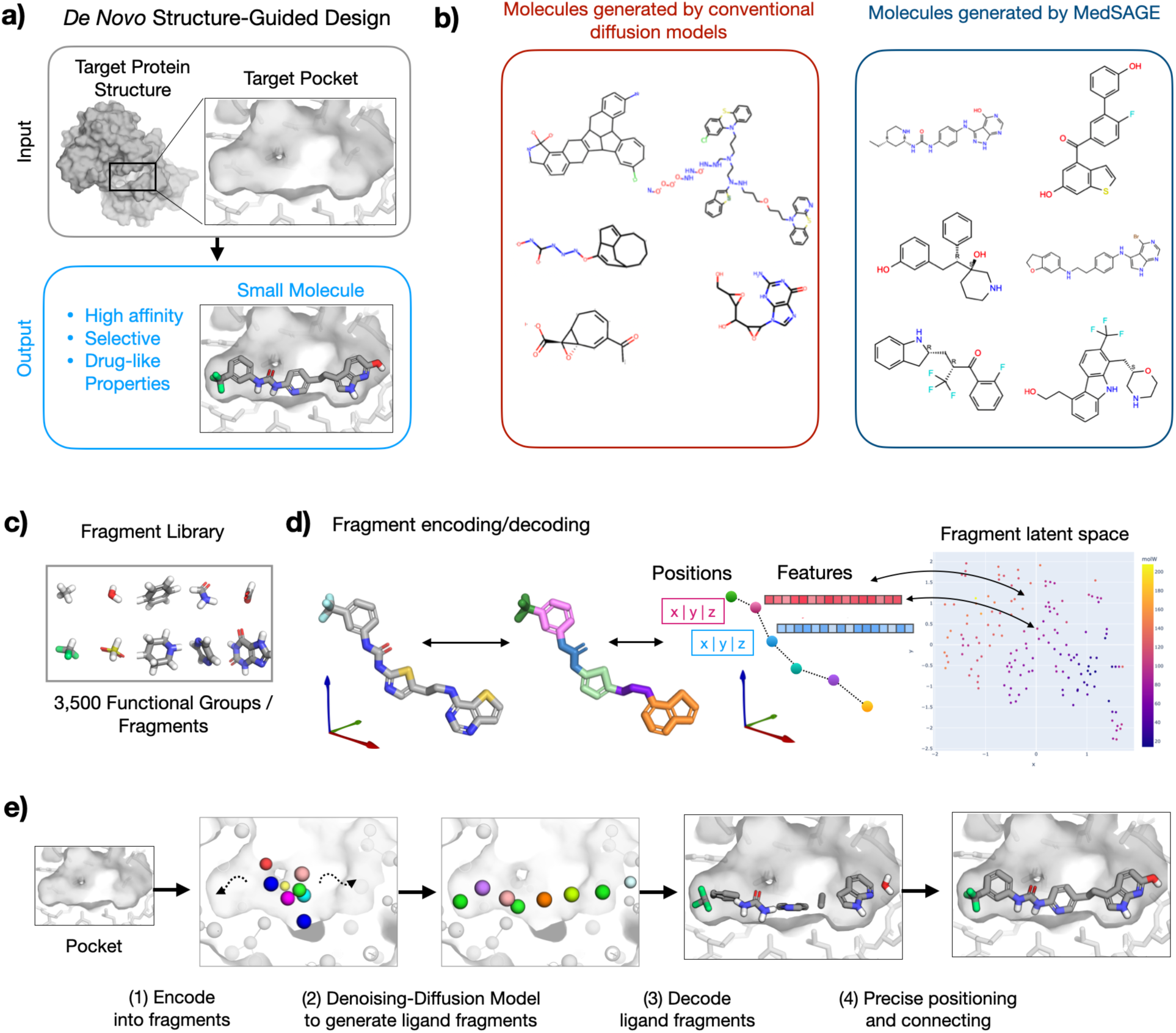
MedSAGE is a generative model for structure-guided design of small molecule drugs that integrates diffusion models with molecular fragments relevant to medicinal chemistry. **(a)** Overview of *de novo* structure-guided drug design: novel molecules are generated using the 3D structure of the target protein. Molecules should be high-affinity, selective, and drug-like. **(b)** Molecules from prior diffusion models (left) often suffer from poor drug-likeness and low synthetic feasibility due in part to complex ring systems and reactive groups (e.g., epoxides). MedSAGE overcomes these issues using a fragment-based diffusion framework (examples of generated molecules shown on right). (c) **Fragment Library**: we curated an extensive set of functional groups and ring systems critical for medicinal chemistry applications. **(d) Fragment Encoding/Decoding**: Molecules are represented as centroids of their constituent fragments, which are labeled by low-dimensional latent vectors for efficient use in diffusion models. The latent space embedding smoothly encodes chemical and geometric properties of the fragments. **(e) MedSAGE Process**: The generative model operates in two phases. The first phase operates on fragments; fragments are positioned and assigned types simultaneously within the context of the target protein binding pocket using a trained denoising diffusion model. In the second phase, the coarse-grained fragment representation is converted back to atomic-level structures, followed by connecting nearby fragments with a custom algorithm.

We hypothesized that diffusion models can be adapted to effectively generate small molecules by reconsidering their fundamental building blocks from a chemistry perspective. Diffusion models have demonstrated significant success across domains where data can be represented as grids or sequences, including image generation^17,18^ (pixels), protein design^5^ (amino acid sequence), and natural language^19^ (word embeddings). However, complex small molecule structures are not naturally represented by simple sequences or grids. 3D point clouds of atoms are a relatively straightforward representation, but medicinal chemists typically conceptualize small molecules in terms of small chemical groups or “fragments” rather than individual atoms and elements; these fragments are the natural vocabulary of drug design^20–24^. However, using fragments for generative AI also entails unique challenges: thousands of chemical fragments exist with highly diverse properties^22,23^, and these fragments can connect in a myriad of complex ways^25^.

We address the key limitations of previous generative models for small-molecule design by introducing MedSAGE, a framework that combines diffusion models for 3D structures with a fragment-based representation. For training these models, we curate and prepare a custom dataset focused specifically on drug-like small molecules. Our approach achieves state-of-the-art results in *de novo*, structure-guided small molecule design—consistently generating synthesizable, target-specific compounds with favorable drug-like physicochemical properties. We validate MedSAGE through detailed comparisons with previous generative methods and real-world virtual screening campaigns, showing that it can reliably generate high-quality molecules for therapeutically relevant targets. These results mark a turning point in making diffusion models practically useful for real-world, structure-guided small molecule drugs.

### Developing a Fragment-Based Molecular Representation for Diffusion Models

To simplify the learning process and eliminate the generation of chemically implausible or synthetically intractable molecules, we developed a novel molecular diffusion approach that encodes both chemical and geometric information into a lower-dimensional latent space, guided by strong chemical intuition. Drawing inspiration from coarse-graining techniques used in molecular dynamics simulations, we represent molecules—including both ligands and protein targets—as fragments. In this context, fragments refer to small functional groups and ring systems that are relevant for medicinal chemistry applications, distinct from the more complex molecular fragments typically used in fragment-based drug discovery^26^. We then designed and trained diffusion models to operate directly on this fragment-based representation.

We first curated a comprehensive library of several thousand fragments relevant to drug discovery (Fig. 1c, Supplementary Figure 1), extracted from drug-like small molecule ligands (see Methods, Fragment Preparation). Each fragment was then assigned a low-dimensional vector label that encodes a smooth embedding of its Extended Three-Dimensional Fingerprint^27^ along with key chemical properties such as charge, polarity, hydrogen bond donors/acceptors, and atom count (Fig. 1d). These embeddings were learned in an unsupervised manner using t-SNE^28^, enabling a unified embedding of all fragments in the library.

Compared to one-hot encoded fragment labels or atomic element labels, these smooth and normalized vector labels are better suited for diffusion models and improve computational efficiency. The fragment-based representation significantly reduces degrees of freedom—by more than tenfold compared to the all-atom representation commonly used in diffusion models for small molecules. For example, generating a benzene ring requires only a single fragment point and its associated label (six degrees of freedom in total), rather than precisely positioning and labeling all six carbon atoms (6×3 spatial coordinates + 6×8 common elements = 66 degrees of freedom). To encode molecular structures in this fragment representation, we developed a custom algorithm that decomposes molecules into their constituent fragments and assigns a labeled point at each fragment’s centroid (see Methods, Fragment Preparation).

MedSAGE designs molecules targeting a protein binding pocket structure through a two-phase process (Fig. 1e). In the first phase, we apply a denoising diffusion probabilistic model (DDPM) to generate the labels and positions of ligand fragments within the 3D pocket environment. DDPMs are a class of generative models that learn to reverse a noise-adding (diffusion) process by gradually refining noisy inputs back into structured outputs. The target pocket structure remains fixed throughout the noising and denoising process and is encoded using the same fragment representation. To distinguish between ligand and pocket points, we incorporate additional context features that indicate whether each point belongs to the ligand or the pocket. Following prior work, our diffusion model employs an equivariant graph neural network (EGNN) for denoising predictions^11,29^. The EGNN architecture effectively leverages geometric information by considering pairwise distances within the fragment point cloud.

To train our diffusion model, we constructed a custom dataset of 3D structures containing small molecule ligands with explicitly drug-like properties bound to protein targets. We began by downloading and processing all ligand-annotated structures from the Protein Data Bank (PDB). The dataset was then filtered to exclude common biomolecules such as lipids, peptides, carbohydrates, and nucleotides, as well as ligands that fell outside established drug-like property ranges (see Methods, Dataset Preparation). This curation process resulted in a dataset of approximately 35,000 protein-ligand complexes. These complexes were further processed to add hydrogens, fill missing sidechains, and determine protonation states. Target pockets were selected using the known ligands. Special care was taken to account for protonation states in both the dataset and our fragment library — a detail crucial for accurately capturing specific chemical features and interactions.

In the second phase of MedSAGE, we convert the coarse-grained fragment representation back into an atomic structure to produce a final designed molecule. First, the generated fragment labels are decoded into specific fragments structures from our library using the learned latent space map (Fig. 1d). However, this step alone does not specify how nearby fragments should be chemically connected. To address this, we developed a search algorithm that identifies the optimal bond connections between fragments. The algorithm generates candidate molecules by adding bonds between adjacent fragments while enforcing chemical validity rules — for example, ensuring that an sp³ carbon forms no more than four bonds (see Methods, Fragment Connecting). We then apply a physics-based scoring function (Glide) to evaluate the candidates and select the highest-scoring structure^30^. While this connection step could eventually be replaced by a second AI model operating on atomic coordinates, this physics approach enforces strong physical constraints and leverage the robust scoring capabilities of existing physics-based docking methods. Importantly, this method remains computationally efficient because the fragments are already fixed and roughly positioned, significantly reducing the search space for likely connections. In practice, evaluating 40 connectivity candidates is often sufficient, after which scores tend to plateau (Supp. Fig. 2).

### A comprehensive benchmark on therapeutically relevant targets demonstrates MedSAGE’s effectiveness for structure-based drug design

To evaluate the performance of MedSAGE in structure-guided drug design, we constructed a rigorous benchmark of protein targets with therapeutic relevance. Specifically, we selected 25 diverse protein-ligand complexes from the test set, each containing a protein target bound to a high-affinity (<1 µM), drug-like small molecule—termed the “reference ligand”— which is either an approved drug, clinical candidate, or optimized preclinical candidate (Supp. Table 1). These targets (and proteins with similar sequences) were not used during model training. The average binding affinity of the reference ligands was 30 nM. For each protein target binding pocket, we generated 400 novel molecules using MedSAGE (Fig. 2,). Each set of generated molecules was then evaluated across a comprehensive panel of 31 physicochemical and pharmacological properties, including logP, molecular charge, number of rotatable bonds, synthetic complexity, and predicted binding affinity (estimated via physics-based scoring functions) (Fig. 3, Supp. Table 2). To place MedSAGE’s performance into context, we conducted parallel evaluations on the same benchmarking targets using several leading generative AI methods, DiffSBDD^8^, IPDiff^10^, and PMDM^9^, which are reported to be state-of-the-art in peer-reviewed publications from the last year. These methods utilize all-atom diffusion models to generate small molecules, allowing for a direct comparison with MedSAGE’s distinct latent fragment-based approach.

**Figure 2:**
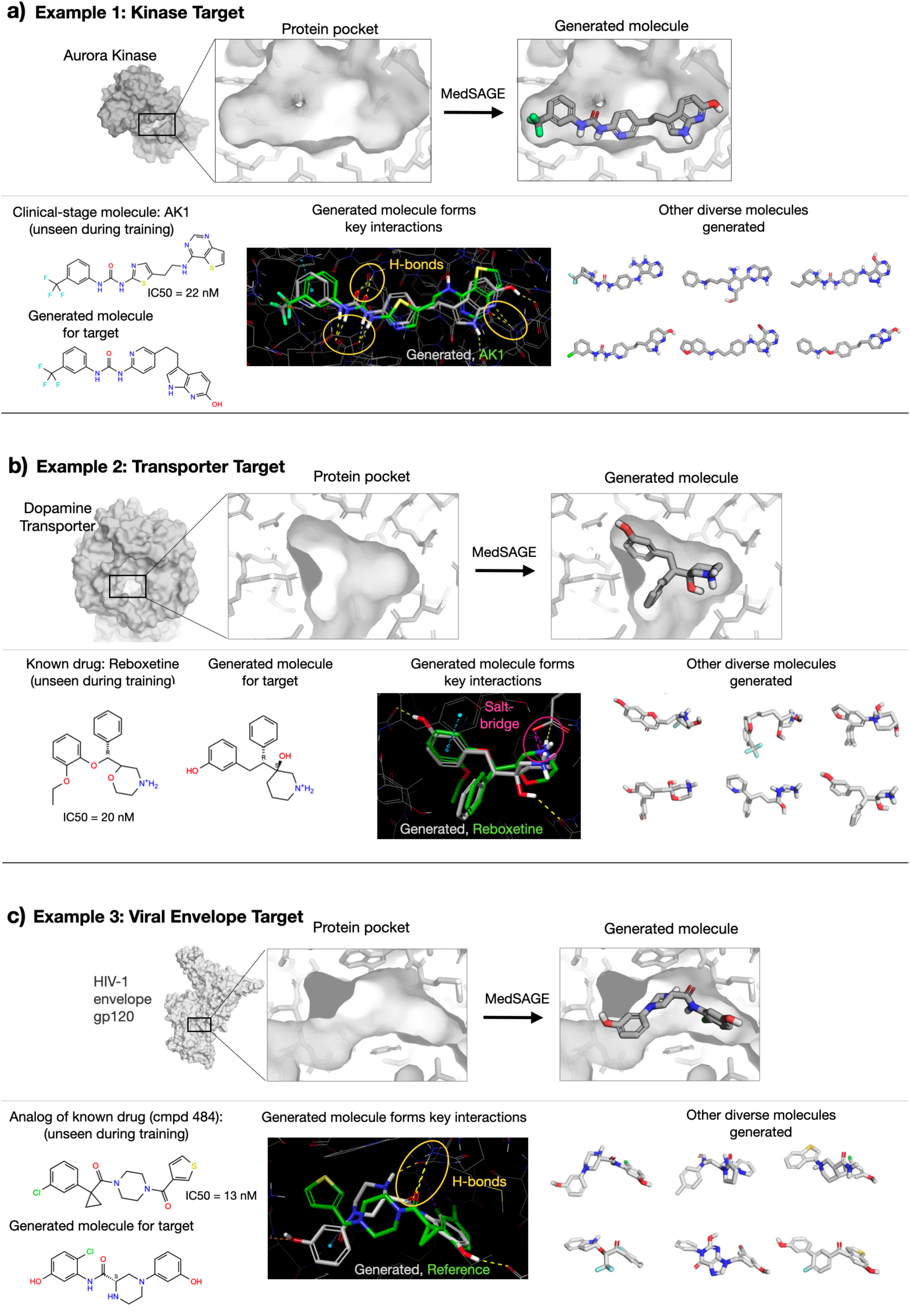
MedSAGE generates realistic, diverse molecules that recapitulate scaffolds and key interactions of high-affinity drugs and clinical candidates. Selected examples of MedSAGE-generated molecules for three drug targets in the test set. For each target, we show the protein pocket structure (light grey surface and sticks) provided to MedSAGE, along with a generated molecule (dark grey sticks). We compare the generated molecule to the “reference ligand”—a drug-like molecule with high affinity for the target protein. Dark-background images illustrate ligand-pocket interactions, including hydrogen bonds (yellow), salt bridges (magenta), and π–π interactions (blue). For each example, we show an additional six MedSAGE-generated molecules to highlight the diversity of generated chemical structures. **(a)** Target is a crystal structure of Aurora kinase (PDB 3D14), a serine/threonine kinase essential for mitosis and implicated in various cancers. The reference ligand is a potent and selective clinical-stage candidate with a binding affinity of 22 nM and demonstrated *in vivo* antitumor activity. **(b)** Target is the dopamine transporter from *D. melanogaster* (PDB 4XNX), used to study the binding mechanisms of antidepressants to various transporters. The reference ligand is reboxetine, an approved antidepressant with 20 nM affinity for this target. **(c)** The target is the gp120 subunit of the HIV-1 envelope trimer (PDB 6MTN). The reference ligand, cmpd 484, is an analog of BMS-626529 (temsavir), an approved antiretroviral medication.

**Figure 3:**
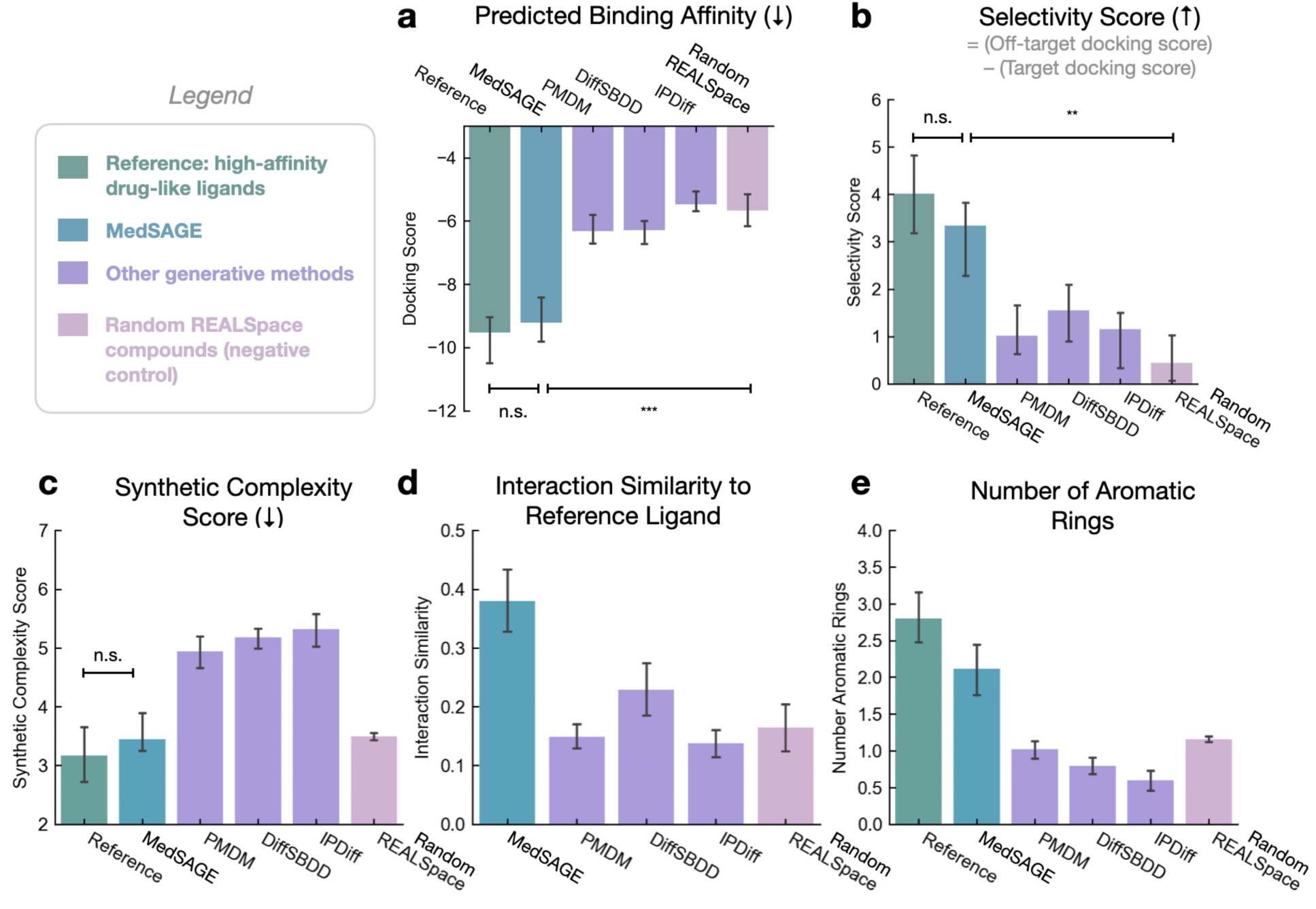
MedSAGE generates molecules with superior affinity and selectivity compared to other diffusion-based methods for small molecule generation. Benchmarking was conducted using a curated dataset of 25 diverse proteins, each of therapeutic importance. For each target, 400 molecules were generated by each method, and their properties were evaluated. The “reference ligand” set consists of known drug-like ligands with high-affinity for the target proteins serving as the positive control. Reference ligands consist of approved drugs, clinical candidates, and optimized preclinical candidates. PMDM, DiffSBDD, and IPDiff are molecules generated using publicly available all-atom diffusion models, each applied to our benchmarking set of proteins. The “Random REALSpace” set consists of molecules randomly sampled from the Enamine REALSpace chemical library, without using the target protein structure, serving as the negative control. Data are presented as the median values, with 95% confidence intervals (see Methods for details on statistical analyses). *P*-values are calculated using two-sided t-test (n.s. indicates p>0.05, * p<0.05, ** p<0.01, *** p<0.001).

Overall, MedSAGE demonstrates substantial improvements over existing leading methods by generating molecules that are highly realistic from a medicinal chemistry perspective, closely matching the structural and physicochemical properties of the reference ligands and adhering to established drug design rules (Fig. 2, Fig. 3, Supp. Tables 2 and 3). This is despite not explicitly incorporating physicochemical property predictions or objectives into the generative process. As further illustrated in the Case Studies below, MedSAGE is able to generate scaffolds strikingly similar to the known reference ligands despite having no prior exposure to them during training (Fig. 2). Additionally, molecules generated by MedSAGE frequently recapitulated key protein-ligand interactions, such as hydrogen bonds observed in the original high-affinity complexes (Fig. 2a). This indicates that MedSAGE effectively captures fundamental principles underlying ligand-protein binding.

### MedSAGE generates molecules with high predicted affinity and selectivity

The first metric we assessed was the predicted binding affinity of the generated molecules, approximated using docking scores from Schrödinger’s Glide (Fig. 3a)^30^. The median docking score of molecules generated by MedSAGE closely matched that of the high-affinity reference ligands (−9.2 vs. −9.5; difference not significant, p > 0.05, t-test; Supp. Table 2). In contrast, molecules generated by other diffusion-based methods had weaker median docking scores, ranging from −5.5 to −6.3 (Fig. 3a, Supp. Table 3).

To further contextualize these results, we introduced a negative control by randomly sampling molecules from Enamine REALSpace and evaluating their docking scores and other properties^31^. These control molecules, not designed or optimized for the protein targets in any way, served as a baseline and negative control. Their median docking score was −5.7 (Fig. 3a), significantly worse than MedSAGE-generated molecules (p < 0.001, t-test) but not statistically different from molecules generated by IPDiff and DiffSBDD (p > 0.05, t-test). This finding suggests that these alternative methods struggled to generate molecules specifically tailored to the target protein pockets.

Beyond assessing docking scores at the target protein, we also introduced a selectivity metric defined as the difference in docking scores between the intended target and a set of off-target proteins randomly selected from the benchmark set. Selectivity is crucial in drug design, as molecules optimized solely for affinity (or docking scores) can become overly bulky or hydrophobic, leading to non-specific binding to off-target proteins. MedSAGE-generated molecules displayed selectivity comparable to the reference ligands (Fig. 3b; 3.6 vs. 4.2, p > 0.05, t-test; Supp. Table 2). In contrast, molecules produced by the all-atom diffusion methods showed substantially lower selectivity (1.3–1.5), only slightly above that of the negative control molecules.

We hypothesized that the better selectivity of MedSAGE-generated molecules could stem from their ability to form specific key interactions—such as hydrogen bonds, salt bridges, and pi–pi stacking interactions—with the target proteins. To evaluate this, we quantified what fraction of interactions made by reference ligands were reproduced by each set of generated ligands. On average, MedSAGE-generated molecules reproduced 35% of the interactions observed in the high-affinity reference ligands versus 16% for the negative control (p<0.001, t-test). In comparison, molecules generated by other diffusion-based methods reproduced only 12– 23% of reference interactions.

### MedSAGE Produces Structurally Diverse, Synthesizable, and Drug-Like Molecules

We next evaluated broader physicochemical properties relevant to synthesizability and crucial for developing drug candidates with favorable absorption, distribution, metabolism, and excretion (ADME) profiles. Across the majority of these metrics, MedSAGE-generated molecules closely matched the properties of the reference ligands (Supp. Fig. 3, Supp. Table 2) MedSAGE also demonstrated the ability to generate structurally diverse ligands, even within individual protein targets. The average pairwise Tanimoto fingerprint similarity was less than 0.1 between pairs of generated ligands for a given target. Additionally, out of 400 ligands generated per target, there was an average of 197 unique scaffolds (see Methods; Benchmarking).

We assessed synthetic feasibility using a standard synthetic complexity score ranging from 1 (easy to synthesize) to 10 (difficult). MedSAGE-generated molecules had low average complexity scores (3.5), similar to reference ligands (3.2; p > 0.05, t-test). In contrast, DiffSBDD, IPDiff, and PMDM produced molecules with significantly higher scores (4.9–5.3), reflecting their use of complex scaffolds and rare chemical groups. All-atom diffusion methods generated molecules with more stereocenters (>3 per molecule) than MedSAGE or REALSpace (1.3 and 1.4, respectively), and a higher frequency of complex ring systems such as bridged or spiro atoms (Supp. Table 3).

MedSAGE produced molecules with distributions of ring counts, rotatable bonds, and fraction of sp³ carbons closely resembling those of the reference ligands. However, we observed that MedSAGE-generated molecules contained slightly fewer heterocyclic rings compared to the reference ligands, perhaps reflecting a tendency to select simpler and more common ring fragments. In contrast, other diffusion-based methods (DiffSBDD and IPDiff) tended to generate molecules with fewer aromatic rings and a far higher prevalence of aliphatic rings (>3 aliphatic rings) than reference ligands (<1 aliphatic ring on average). This discrepancy likely arises from challenges in precisely positioning atoms during the diffusion generation process to form planar aromatic rings. MedSAGE avoids this issue because it generates molecules directly from fragment space, where different ring systems have distinct properties and are explicitly differentiated.

MedSAGE-generated molecules exhibited physicochemical properties closely aligned with medicinal chemistry guidelines, such as the Lipinski’s Rule of 5; these properties include molecular weight (340 Da, recommended <500 Da), number of hydrogen bond donors (3.0, recommended <5), hydrogen bond acceptors (4.2, recommended <10). The average logP was 2.0, placing it within the ideal range (0–5) for oral bioavailability and CNS penetration, which is often desirable for therapeutics. In contrast, molecules generated by IPDiff and DiffSBDD tended to be more polar, with lower logP values of 0.87 and 0.7, respectively. Overall, these findings suggest that MedSAGE effectively generates molecules with physicochemical properties aligned with established principles of drug design.

## Case Studies

We next evaluated MedSAGE in several detailed case studies (Figure 2). In these examples, MedSAGE demonstrated a remarkable ability to “rediscover” scaffolds similar the reference ligands—despite not being trained on these known actives. Given the model’s ability to generate trillions of possible molecules, such convergence is highly unlikely to occur by chance and suggests that MedSAGE captures key medicinal chemistry principles underlying high-affinity, drug-like ligand design.

In the first case study, the target is Aurora kinase, a serine/threonine kinase that plays a critical role in mitosis and is implicated in various cancers. The reference ligand, AK1 (also known as SNS-314), is a clinical-stage candidate that acts as a potent and selective Aurora kinase inhibitor with a binding affinity of 22 nM, and significant activity in tumor models. With the Aurora kinase pocket as input, MedSAGE generated a molecule with a scaffold similar—but not identical—to AK1 (Figure 2a). The generated compound preserves key structural features, including a core urea group flanked by two aromatic rings. This urea group forms critical hydrogen bonds with an aspartate and lysine residue in the binding pocket. Additionally, both the reference and MedSAGE-generated molecules feature a heterocyclic aromatic ring at the end of the pocket that forms hydrogen bonds with a backbone amide. The docking score of the MedSAGE-generated compound is -12, reflecting strong predicted binding affinity, with an off-target average docking score of -5, indicating high selectivity for this target over off-target pockets.

In the second case study, the target is a dopamine transporter (DAT) bound to reboxetine, an approved antidepressant with 20 nM affinity for this target protein. MedSAGE generated a molecule with a scaffold closely resembling that of reboxetine, and a similar structural overlay (Fig. 2b). Two aromatic rings insert into a hydrophobic cleft, and a secondary amine within an aliphatic ring forms a salt bridge with a conserved aspartate residue. Notably, the generated molecule forms two additional hydrogen bonds compared to reboxetine and has a lower docking score (−9.7 vs. -8.4 for reboxetine).

While this particular molecule shares a similar scaffold with reboxetine, other MedSAGE-generated compounds display a wide range of diverse chemical structures (Fig. 2b). Nevertheless, all retain a key amine functional group that forms the characteristic ionic interaction with the aspartate—a hallmark of nearly all DAT and NET (norepinephrine reuptake) inhibitors. In other words, MedSAGE consistently captures the essential features required for binding while sampling diverse scaffolds that embody those features.

In our final case study, the target is the gp120 subunit of the HIV-1 envelope (Env) trimer (Fig. 3c). The reference ligand, compound 484, is an analog of Temsavir (BMS-626529), an approved antiretroviral medication. Like compound 484 and Temsavir, the MedSAGE-generated molecule contains a central piperazine ring, a carbonyl group that forms a key hydrogen bond, and a phenyl ring positioned in a narrow pocket—key structural features shared across the class of known inhibitors. Notably, the generated molecule forms a hydrogen bond with an aspartate residue that is not made by compound 484, but is made by temsavir, which is known to be more potent. The piperazine ring adopts a different geometry in the generated molecule, enabling an additional hydrogen bond interaction.

Interestingly, across these three case studies, we observe that MedSAGE-generated molecules tend to form more hydrogen bonds with the target protein than the reference ligands. This is also reflected in the higher average number of hydrogen bond donors (Supp. Fig. 3). While these interactions may enhance binding affinity and selectivity in some cases, they could also reduce membrane permeability, oral absorption, and blood-brain barrier crossing. These design trade-offs will be important to consider in future work.

### MedSAGE identifies high-scoring ligands more efficiently than traditional virtual screening

Virtual screening (VS) is a widely used computational approach for identifying drug candidates by evaluating large chemical libraries with physics-based or AI scoring functions^31,32^. However, the exponential increase in the size of these libraries—now reaching hundreds of billions to trillions of compounds—has dramatically increased computational demands. Generative methods offer a potential approach to complement or even replace traditional VS; method such as MedSAGE could directly generate chemically diverse molecules with high predicted affinity.

To assess the potential of generative methods for this task, we compared MedSAGE to VS across three case studies using retrospective datasets from our laboratory. These datasets correspond to real world VS projects for three distinct drug targets: a transcription factor and two G protein-coupled receptors (GPCRs). The selected targets span a range of binding pocket characteristics, from a shallow interface to a deep, buried pocket. The VS datasets were collected by docking and scoring tens to hundreds of millions of ligands from the Enamine REALSpace using Schrodinger’s Glide (>50 weeks of compute time per target). MedSAGE was used to generate 2000 molecules at each target.

MedSAGE enriched diverse, high-scoring molecules more efficiently than virtual screening (VS) across all three case studies (Figure 4a). The median docking score of the top 10% of MedSAGE-generated molecules was better than that of the top 0.1% of compounds identified through VS. This trend increased at higher percentiles, where the median docking score of the top 1% of MedSAGE-generated molecules was comparable to that of the top 0.001% of VS hits (equivalent to selecting the top 10 out of every 1 million docked compounds).

**Figure 4.**
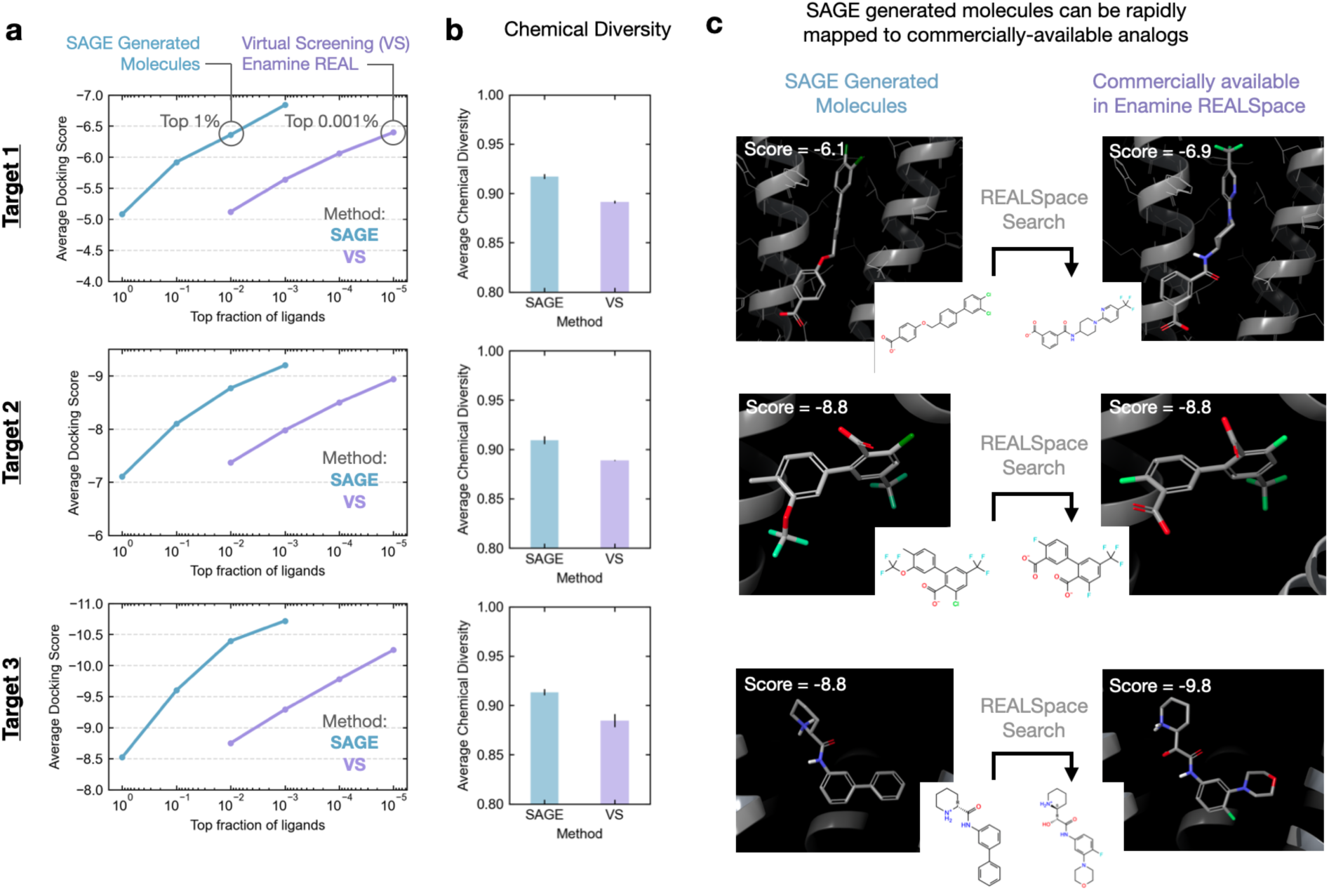
Comparing MedSAGE to traditional virtual screening. **(a)** MedSAGE efficiently identifies high-scoring ligands compared to traditional virtual screening (VS) of chemical libraries. Docking score distributions are shown for three drug targets, with molecules identified through VS (purple lines) and molecules generated by MedSAGE (blue lines). The median Glide docking score is plotted as a function of increasing percentile thresholds. The chemical libraries docked for VS contained 30 million molecules for targets 1 and 2 and 400 million for target 3. MedSAGE was used to generate 2000 molecules for each target. **(b)** Chemical diversity of top-scoring MedSAGE-generated molecules compared to VS-identified molecules, measured by 1–TC (Tanimoto Coefficient) of pairs of molecular fingerprints. Higher values indicate greater diversity. This analysis used the top-scoring 1000 molecules from each method. **(c)** Images at left show selected examples of top MedSAGE-generated ligands (top 1000). These molecules were matched to structurally similar, commercially available compounds from the Enamine REALSpace using efficient analog search algorithms. Docking results show that the matched compounds (shown at right) retain similar predicted binding scores to the original MedSAGE molecules.

MedSAGE-generated molecules were also chemically diverse, with an average Tanimoto coefficient of less than 0.1 between pairs of molecular fingerprints, indicating low structural similarity. The top 1k molecules from VS showed slightly lower diversity, with an average Tanimoto coefficient just above 0.1. Similarly, the top 1,000 ligands contained 595 unique scaffolds on average for MedSAGE and 560 for VS. The findings were consistent across all three drug targets, which reflected a wide range of docking scores.

A potential limitation of MedSAGE is that its generated molecules may be less readily synthesizable than those from combinatorial libraries, which are typically composed of simple, predefined building blocks and reactions. One promising strategy to address this is to first use MedSAGE to rapidly generate promising candidates, and then search for commercially available analogs within chemical libraries using efficient similarity-based algorithms^33,34^. To evaluate this approach, we demonstrated on several examples that MedSAGE-generated molecules were matched to chemically similar library compounds that also achieve favorable docking scores. This workflow offers a scalable path to rapidly identify high-scoring, commercially accessible molecules in trillion compounds libraries.

## Discussion

In this work, we developed MedSAGE, a state-of-the-art generative model for the *de novo* design of small molecules targeting protein structures, capable of rapidly designing drug-like candidates with high predicted affinity. While diffusion models have achieved remarkable success in generating images and even protein structurers, their application to small-molecule design has often resulted in molecules that are often unfeasible from a chemistry perspective. We reimagine diffusion-based methods for medicinal chemistry, using functional groups—rather than individual atoms—as building blocks and mathematically encoding them through smooth embeddings of chemical space. This representation, which drastically reduces the degrees of freedom, is particularly advantageous given the relatively small amount of high-quality training data—fewer than 40,000 examples. Our two-phase workflow—first placing fragments in 3D via a diffusion model and then establishing chemical bonds under physics-based and chemical validity constraints—merges the strengths of AI-driven generation with chemistry intuition.

We conducted a rigorous and comprehensive evaluation of MedSAGE alongside other leading methods, benchmarking performance across a diverse set of protein targets and metrics. MedSAGE consistently achieved state-of-the-art results, generating molecules with docking scores on par with high-affinity approved drugs and clinical candidates. Beyond binding affinity, the method produces ligands with structural and physicochemical properties that align with medicinal chemistry best practices. Notably, these outcomes emerge without explicitly optimizing for such properties during generation—suggesting that generative AI, when structured correctly, can implicitly learn and apply medicinal chemistry principles that are otherwise challenging to model.

Although MedSAGE demonstrates promising performance, there are several important caveats. Currently, MedSAGE does not account for protein flexibility or conformational dynamics. Protein structures used for training and benchmarking were solved in complex with ligands (with a diverse range of affinities). This setup reflects a common real-world use case, as holo structures (ligand-protein complexes) are a common starting point for virtual screening. In some settings, however, available structures may be apo (ligand-free) or derived from computational predictions, which may present less optimal binding pocket conformations.

Furthermore, evaluation of MedSAGE-generated molecules currently depends on physics-based scoring functions, such as Glide, which, although industry standard in computational drug discovery, can introduce biases and overly optimistic estimates of binding affinity. While generated molecules perform well across conventional benchmarks, satisfying these metrics alone may not fully capture molecular quality necessary to develop real drugs. This underscores the need for more robust and sophisticated evaluation metrics, analogous to those used to evaluate realism in generative image models.

MedSAGE unlocks numerous opportunities for future research directions. One potential opportunity lies in developing fine-tuned models for specialized ligand classes—such as macrocycles, covalent inhibitors, and molecular glues—that have historically been challenging for physics-based and structure-based modeling. Additionally, integrating generative models for small molecules with models for protein structure opens new possibilities to capture protein dynamics and flexibility. Unlike conventional docking methods, which suffer from exponential computational costs when modeling protein dynamics, diffusion-based models like MedSAGE can seamlessly incorporate protein flexibility with minimal additional inference cost. In summary, MedSAGE represents a significant advancement toward more precise, adaptable, and data-driven approaches in structure-based drug discovery.

## Contributions

A.S.P. and R.O.D. conceived the project. A.S.P. and T.L. developed embedding method. R.K., M.K., M.X. and S.C.G. assisted with benchmarking. A.S.P and R.O.D. wrote the paper with input from all authors.

## Acknowledgments

Funding was provided by National Science Foundation Graduate Research Fellowships (A.S.P.) and National Institutes of Health (NIH) grant R01GM127359 (R.O.D.).

## Data and Code Availability

Data and code will be made available after peer-reviewed publication.

## Supplementary Methods

### Fragment Preparation

We created a custom fragment library by combining curated fragments relevant to drug discovery with automatically detected fragments from the prepared ligand dataset (Supplementary Figure 1). We also explicitly accounted for differing protonation and tautomeric states of each fragments (a protonated version of a fragment is a separate fragment in our library). The dataset consists of 3700 unique fragments; however, the vast majority of these occurred very rarely in our dataset of ligands.

To obtain the fragment library, we began by defining a minimal set of functional groups, *{F}*, that are important for drug-like ligands (methyl, ethyl, phenyl, carboxy, nitro, cyclohexyl, sulfonamide etc.). We then iterate over each molecule in our dataset to find additional fragments not in *{F}*. First each molecule is converted to a graph representation. Then we remove subgraphs that match a graph in *{F}* as long as these subgraphs are not part of rings and only connected to other subgraphs by single bonds. Then we split apart any ring structures that are connected by single bonds. The remaining connected components of the molecule graph are considered the non-library fragments and added to *{F}*. We do this for each ligand in our dataset to obtain the complete library. In many cases, fragments can be matched in multiple way. We start with fragments that are *terminal*, they are connected to the rest of the molecule by a single bond. If there are multiple terminal fragment matches, we remove the largest one first.

To create the fragment latent space, we computed a vector of properties for each fragment. This consisted of the Extended Three-Dimensional Fingerprint (Axen et. al. 2017) as well as chemical properties of charge, molecular weight, number of hydrogen bond donors/acceptors, and number of rings. We then used t-SNE (sci-kit learn python library), to embed these vectors into a latent space with three dimensions, which was then normalized to be centered at 0 and have a standard deviation of 1 along each dimension. To separate fragment with labels in the latent space that were too similar (within a distance threshold), we iteratively added a relatively small amount of noise to these labels until all labels were sufficiently separated. We found that this approach gave a smooth and interpretable latent space, where fragments with similar properties were clustered together.

### Dataset Preparation and Splitting

For training and testing datasets, we curated a custom dataset of 3D structures containing small molecule ligands with explicitly drug-like properties. An initial set of ligand-protein complexes were obtained by downloading all structures in Protein DataBank (PDB) containing ligand entities. The dataset was filtered to remove common biomolecules (lipids, peptides, carbohydrates, and nucleotides), frequently occurring ligands, and ligands outside established drug-like property ranges (rotatable bonds and hydrogen bond donors). This resulted in a custom dataset of ∼35,000 protein-ligand complexes.

We then prepared the structures using Schrodinger (v2019-2); missing sidechains were added, bond orders were determined, hydrogens were added, and protonation states were determined using Epik at pH 7.0. To improve computational efficiency, we only included amino acid residues in each binding pocket. To prepare the pocket structures, we selected residues in close proximity to the ligand. However, we also added noise to this pocket selection process to avoid the possibility of revealing information to the model about the exact positions of the ground truth ligand atoms.

We divided both datasets into training (85%), validation (10%), and test (5%) sets, ensuring that no protein in one set shared more than 30% sequence identity with any protein in the other sets. This approach was designed to assess generalizability across diverse proteins. It is important to note that we did not explicitly control for pocket similarity. Designing ligands for similar pockets (e.g., related GPCRs) based on knowledge of one pocket is both a common and important task in practice. However, this limitation should be considered when generalizing to completely unknown pockets.

### Architecture and Training

To position and assign fragments, we used Diffusion Denoising Probabilistic Models (DDPMs). We took inspiration from previous all-atom approaches (Hoogeboom et. al. 2022 and Igashov et. al. 2023) and adapted these approaches to our fragment-based approach.

DDPMs are a class of generative models that learn to reverse a diffusion process by gradually removing noise from a sample. The diffusion process starts with the fragment point cloud and progressively adds noise to the positions and features, eventually transforming them into Gaussian noise. The generative denoising process aims to learn the reverse mapping, starting from the Gaussian noise and iteratively removing the noise to reconstruct the original fragment latent points. During the noising and denoising processes, the pocket fragments remain fixed, and the example is centered at the centroid of the pocket fragments. We add additional context features to each point to specify whether the point belongs to the pocket or to the ligand. The learnable function that models the dynamics of the diffusion process is implemented as a E(3)-equivariant graph neural network (EGNN) following Hoogeboom et. al. where the fragment point cloud is converted to a fully connected graph, with edges labeled with distances between the points.

### Fragment Connecting

Given the 3D arrangement of fragments, we developed a custom algorithm to find fragment connections that result in a single fully connected molecule (detailed in *Supplementary Methods*). This algorithm is partially inspired by junction tree methods previously used for 2D graph-based molecule generation^35^.

The algorithm starts by identifying the “neighbor” relationships between the chemical fragments based on their 3D proximity. It creates a “neighbor graph” where nodes represent fragments, and edges represent potential connections between nearby fragments within a threshold distance. The algorithm then enumerates spanning trees from the neighbor graph. A spanning tree is a subset of edges from the neighbor graph that connects all nodes without forming any cycles. These spanning trees represent possible ways to connect the fragments while ensuring that each fragment is connected to at least one other fragment.

Given a proposed connectivity between the fragments, the algorithm identifies the unique attachment points (positions where fragments can be connected) on each fragment. The algorithm then enumerates all valid way to connect the fragments by considering the available attachment points and simple chemical rules for new bonds (e.g. don’t connect fluorine and oxygen).

The algorithm outputs a list of candidate molecules composed of the set of initial fragments linked in various ways that fit the 3D arrangement. The candidate molecules are then scored using a physics-based scoring function (Glide) in the context of the protein binding pocket. The best-scoring candidate from each unique set of fragments is selected based on the docking score. For large molecules, the connectivity algorithm can generate a large number of candidates for the many ways to link the fragment together. To maintain efficiency, these candidates are randomly subsampled before proceeding to the scoring step. In practice, we find that 30-40 candidates are often sufficient to optimize the connectivity, and scores subsequently plateau (Supplementary Figure 2).

### Benchmarking

#### Number of Fragments

Number of fragments for benchmarking was selected based on the number of fragments in the reference ligand. This same information was provided to all tested diffusion methods. This controlled for size variability. In the future, the number of fragments could be determined by a user selected range, or using a second model to predict the number of fragments or a distribution of the number of fragments.

#### Statistical Analyses

In Figure 3 bar plots, data are reported as median values with 95% confidence intervals. For each of the 25 target proteins, metrics were first averaged across the generated molecules, and the median of these averages was then computed across all target proteins. Therefore, N=25 for statistical analyses in Figure 3; this ensures that molecules generated for a particular target protein are not treated as statistically independent samples.

#### Diversity

For calculating unique scaffolds, we (1) select the Murcko scaffold using rdkit, (2) any hetero atoms in aromatic rings of these scaffolds are substituted with carbon (to avoid trivial substitutions such as benzene vs. pyridine), (3) compare equality of SMILES strings of the scaffold.

## Supplemental Discussion

### Protein Flexibility

All protein structures used for training and benchmarking were solved in complex with high-affinity ligands. Therefore, MedSAGE does not attempt to account for protein flexibility and induced fit binding in the current version. This setup reflects a common real-world use case: identifying or generating candidate ligands for targets with a known binding conformation, which is often the starting point for virtual screening and early drug discovery efforts.

## Supplemental Tables

**Supplementary Table 1:**
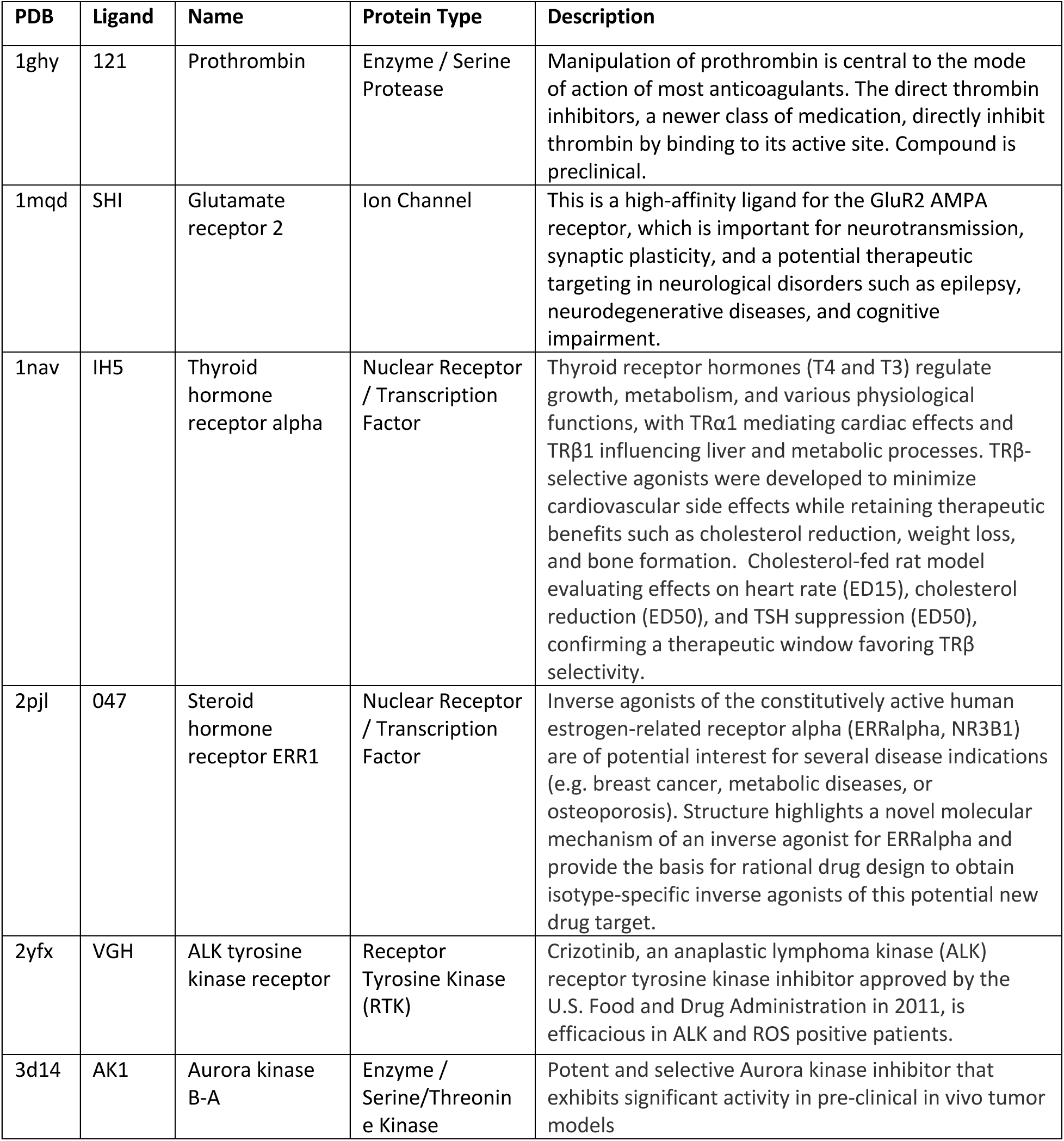
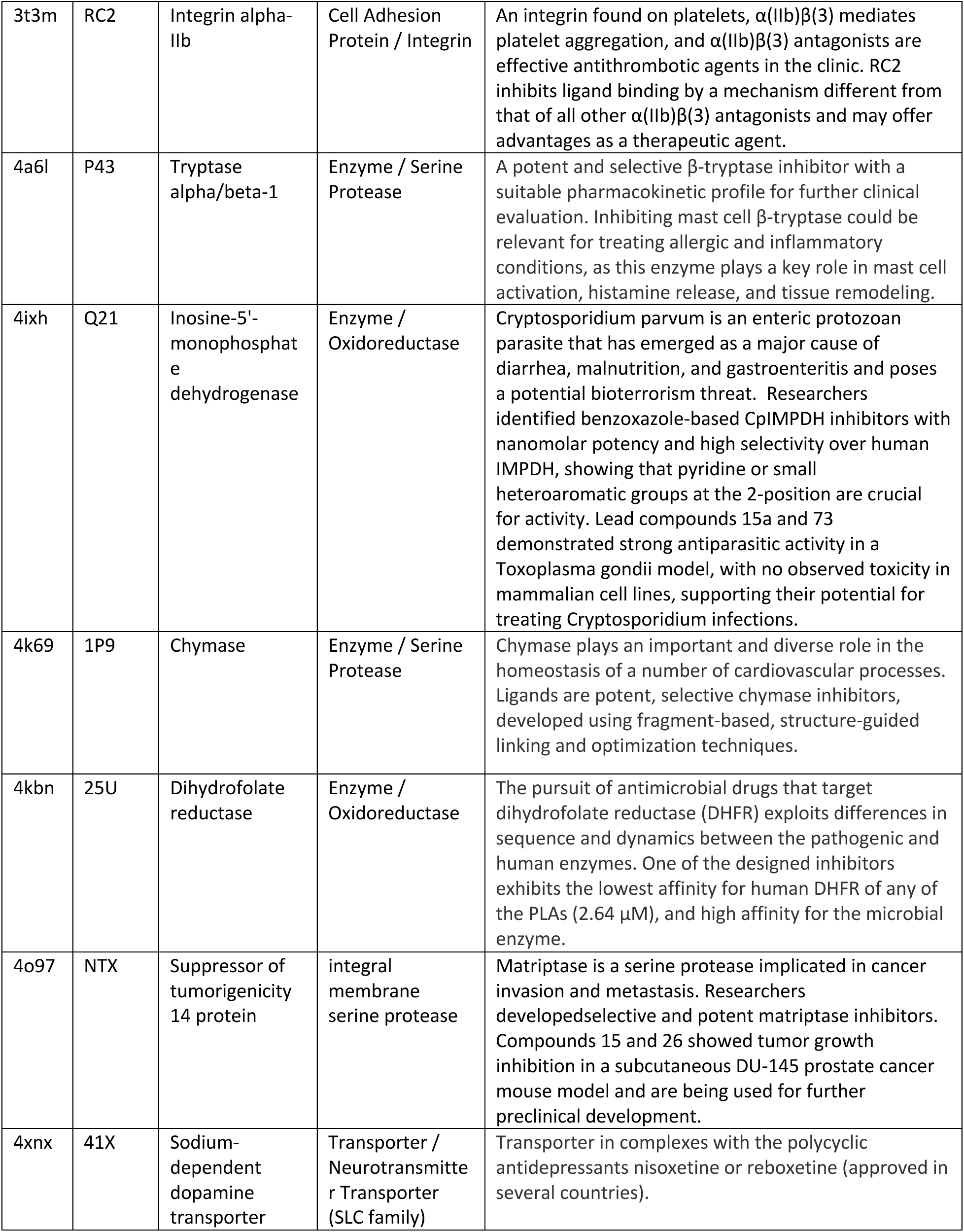
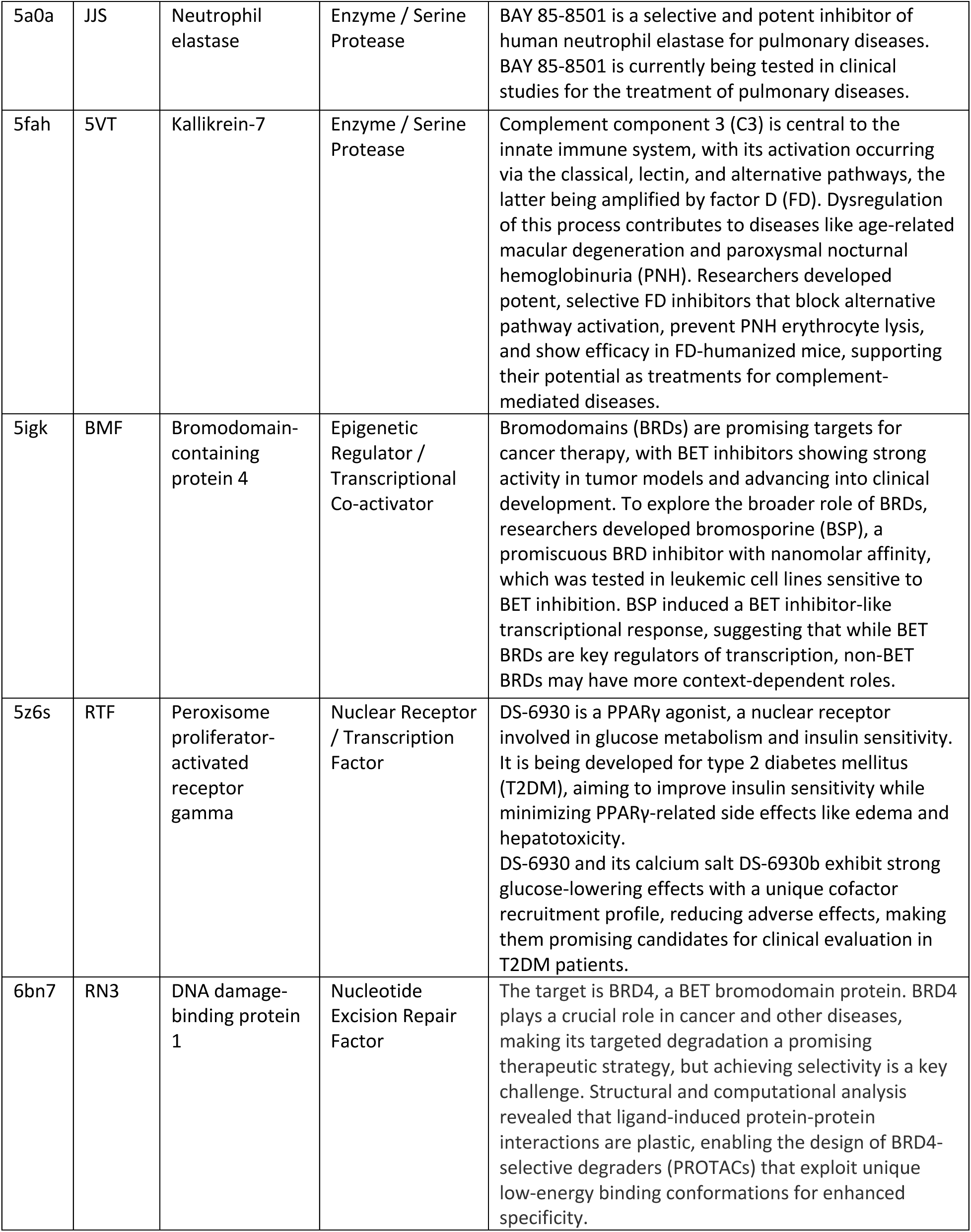
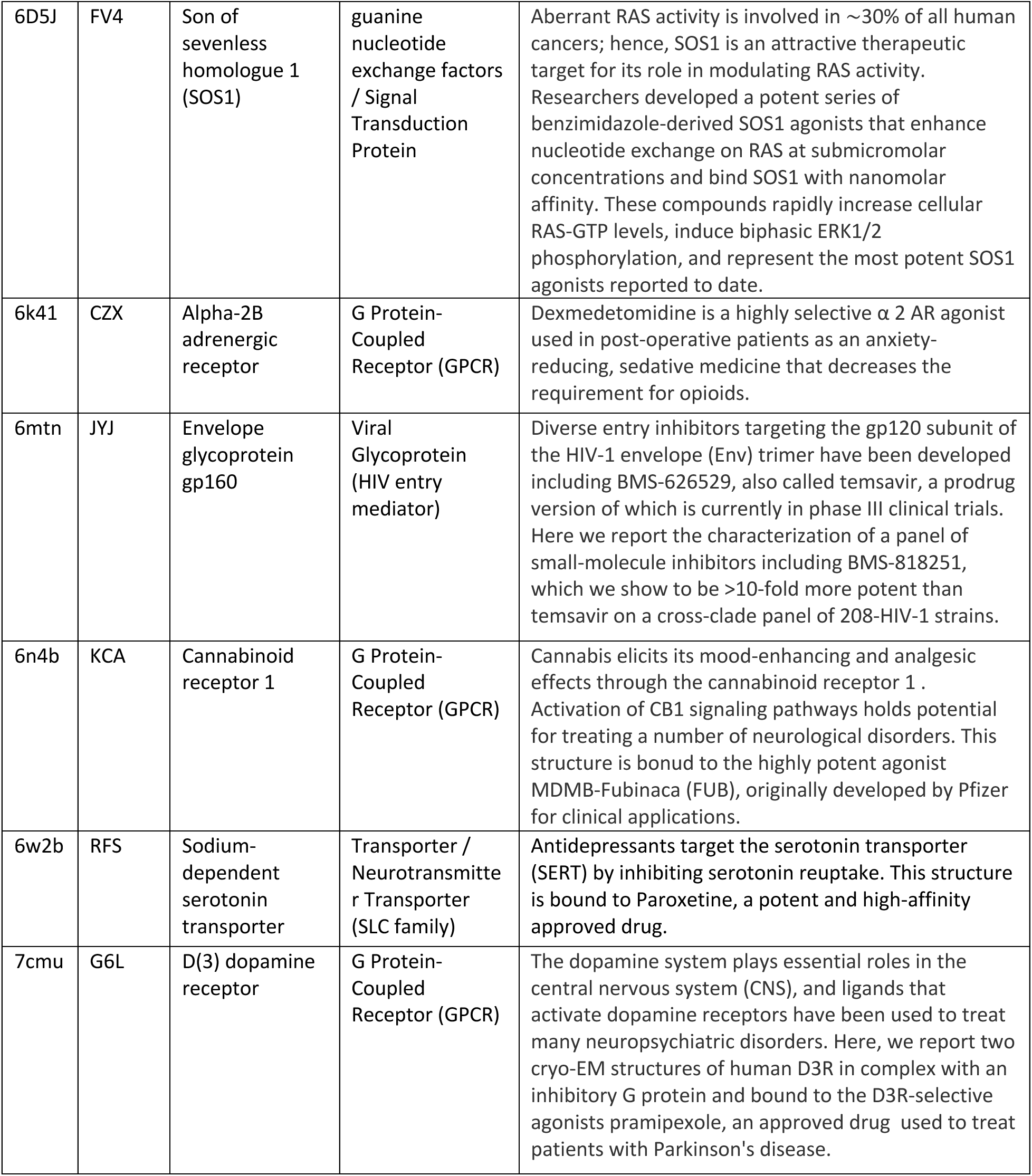
Proteins with Therapeutic Relevance Selected for Benchmarking

**Supplementary Table 2:**
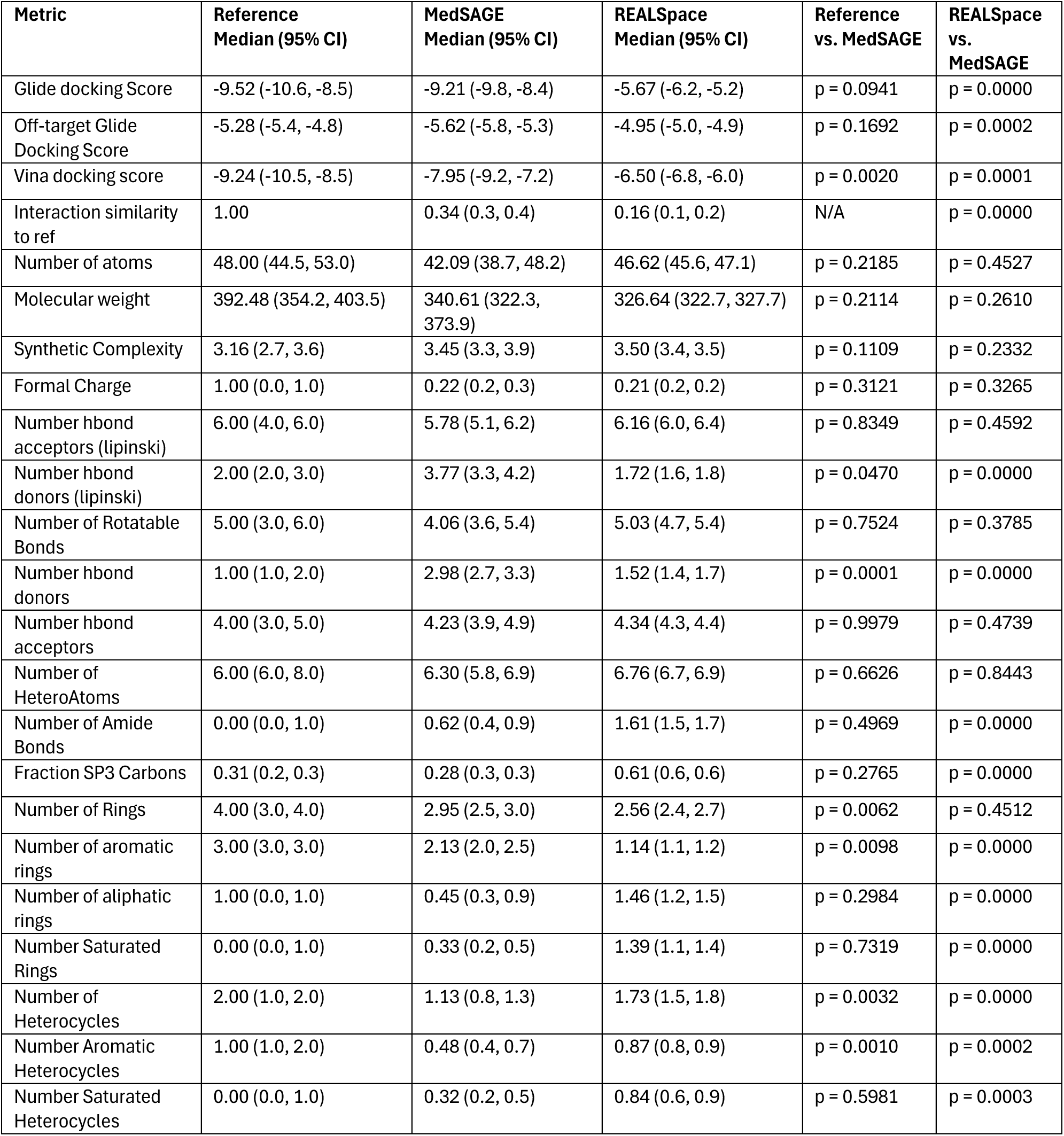
Benchmarking Metrics and Statistics for MedSAGE vs. Reference Ligands vs. REALSpace Metrics are reported as the median with a 95% confidence interval (CI). The Reference vs. MedSAGE column shows p-values from a two-sided Welch’s t-test comparing Reference Ligands to MedSAGE-generated ligands. Similarly, the REALSpace vs. MedSAGE column presents p-values from a two-sided Welch’s t-test comparing REALSpace Ligands to MedSAGE-generated ligands. All statistical calculations were performed by first averaging results for each independent benchmarking protein, then calculating the statistic. Therefore, N = 25 for each metric and comparison.

**Supplementary Table 2:**
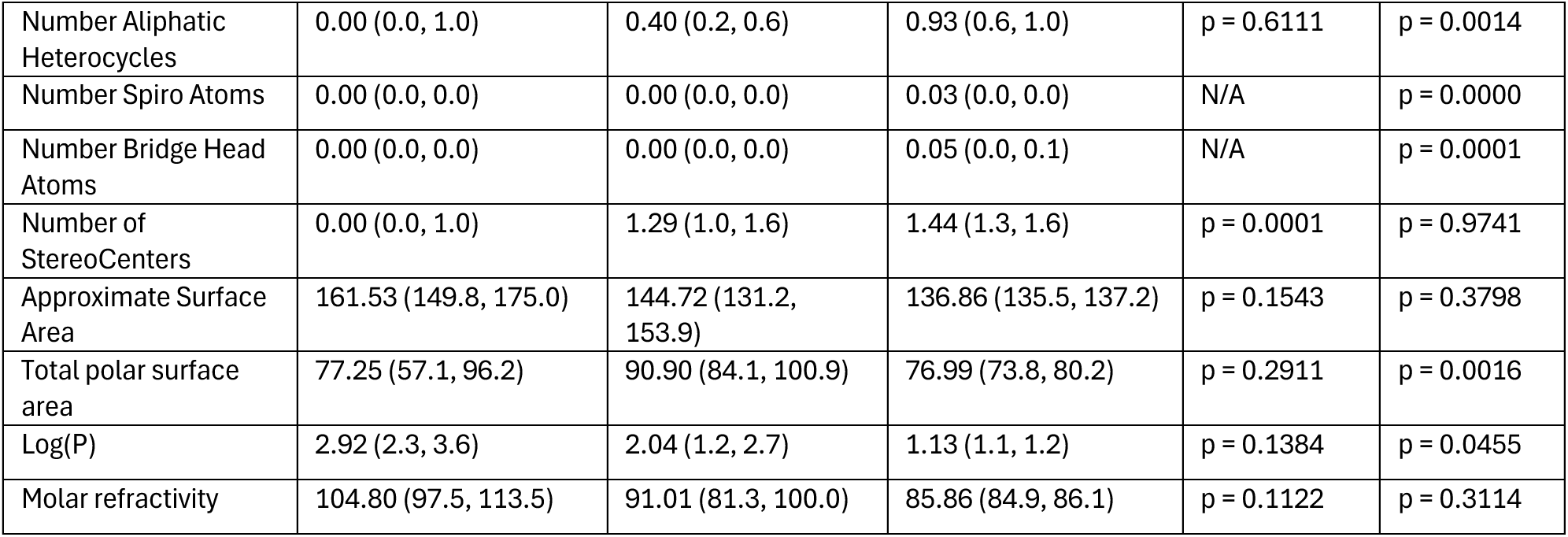
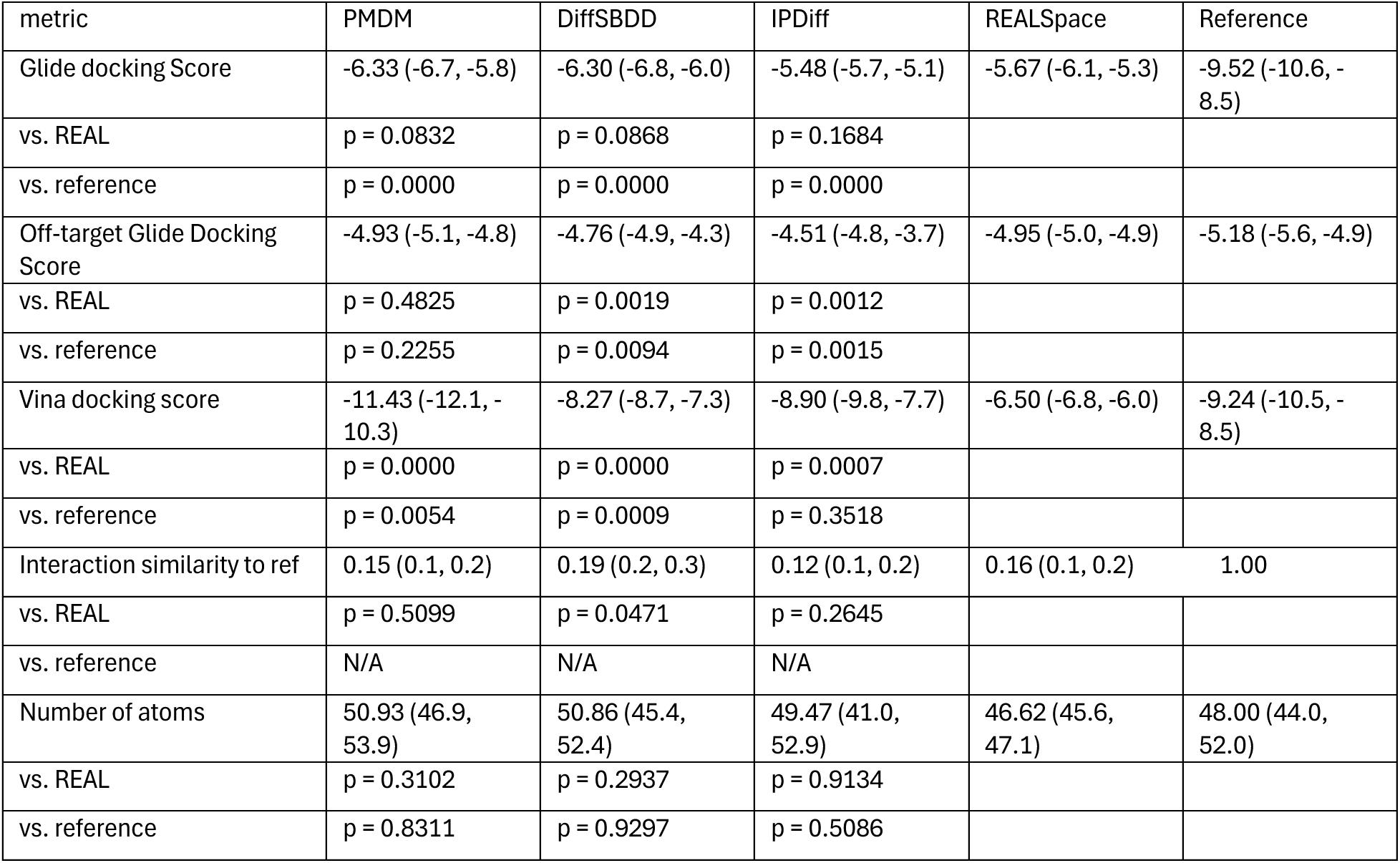
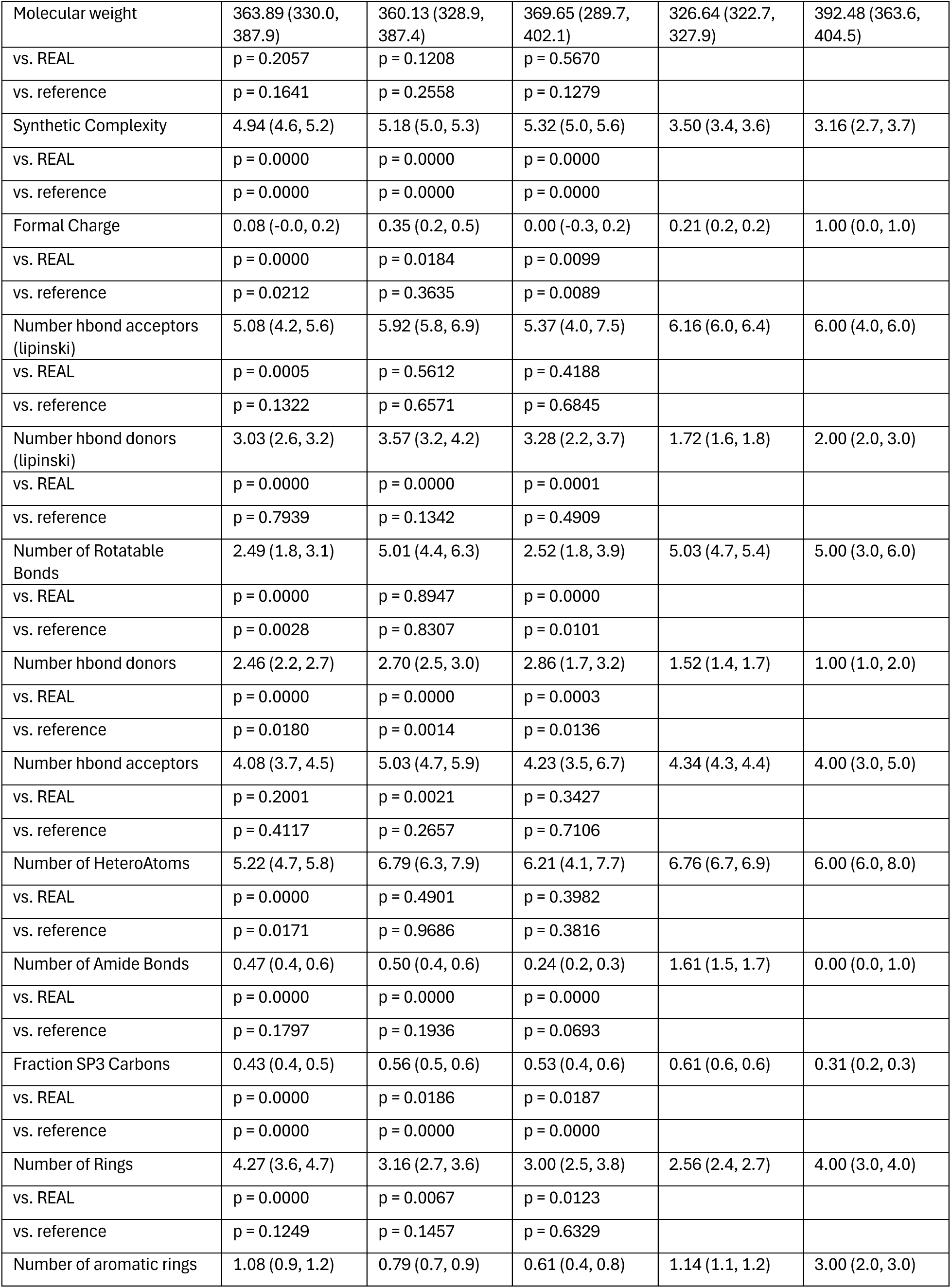
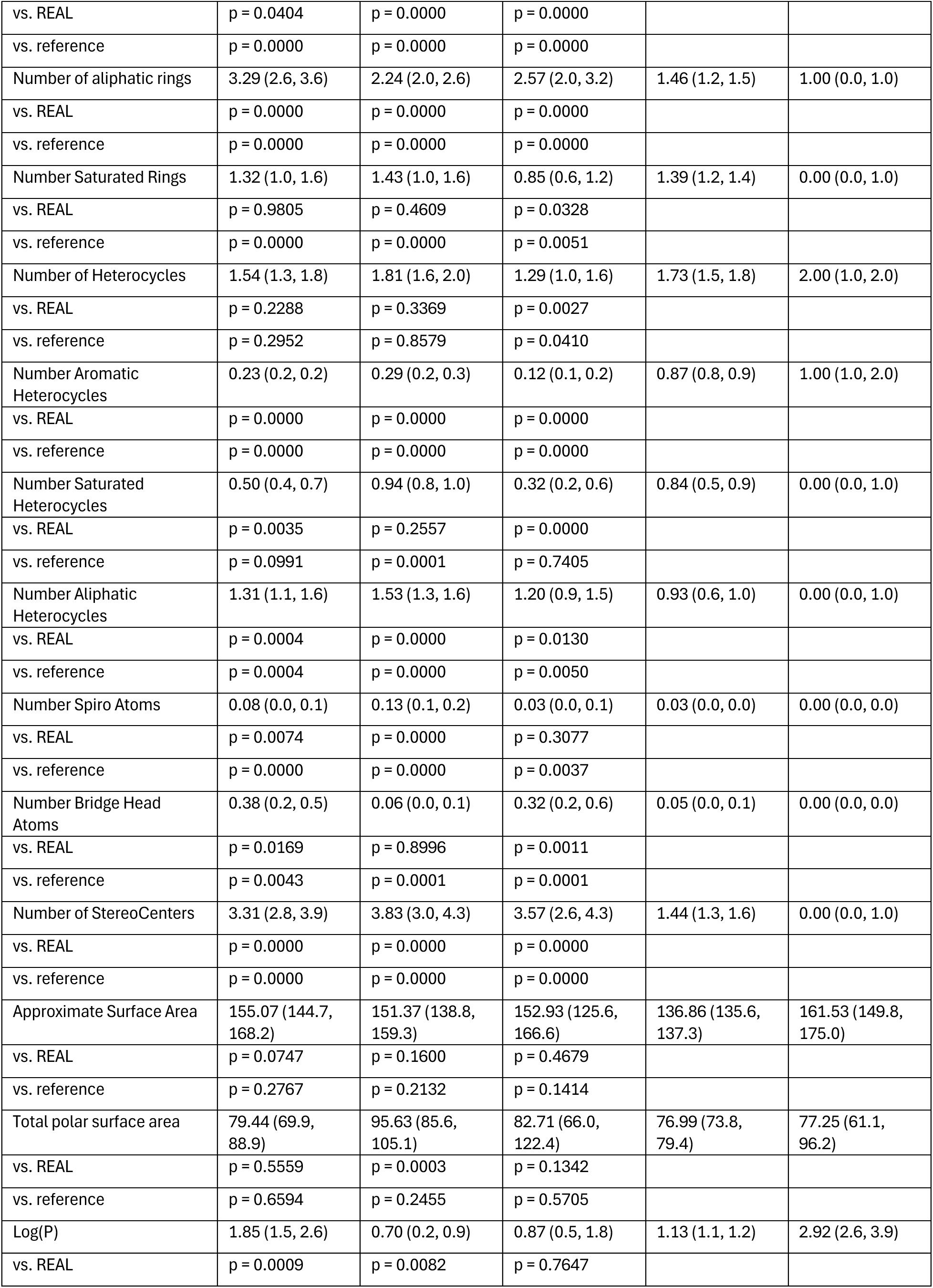
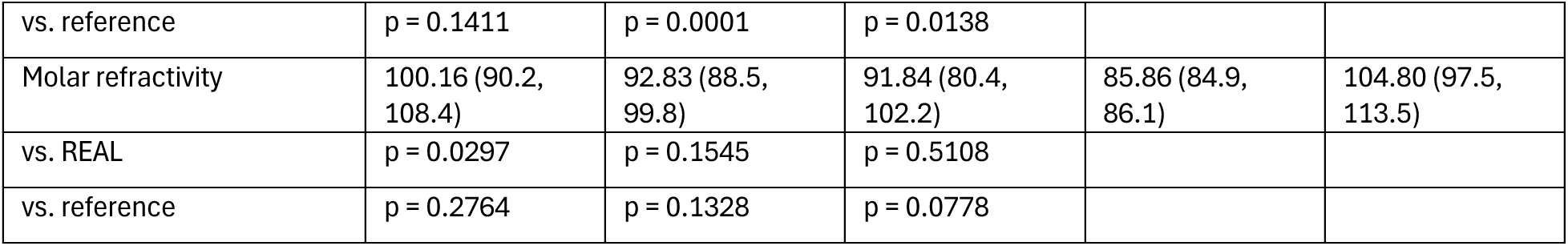
Benchmarking Metrics and Statistics for PMDM, DIffSBDD, IPDiff vs. Reference Ligands and REALSpace Metrics are reported as the median with a 95% confidence interval (CI). The vs. REAL and vs. reference rows presents p-values from a two-sided Welch’s t-test comparing ligands to REALSpace ligands or reference ligands for the metric in the row directly above. All statistical calculations were performed by first aggregating results for each independent benchmarking protein, then calculating the statistic. Therefore, N = 25 for each metric and comparison.

**Supplementary Figure 1.**
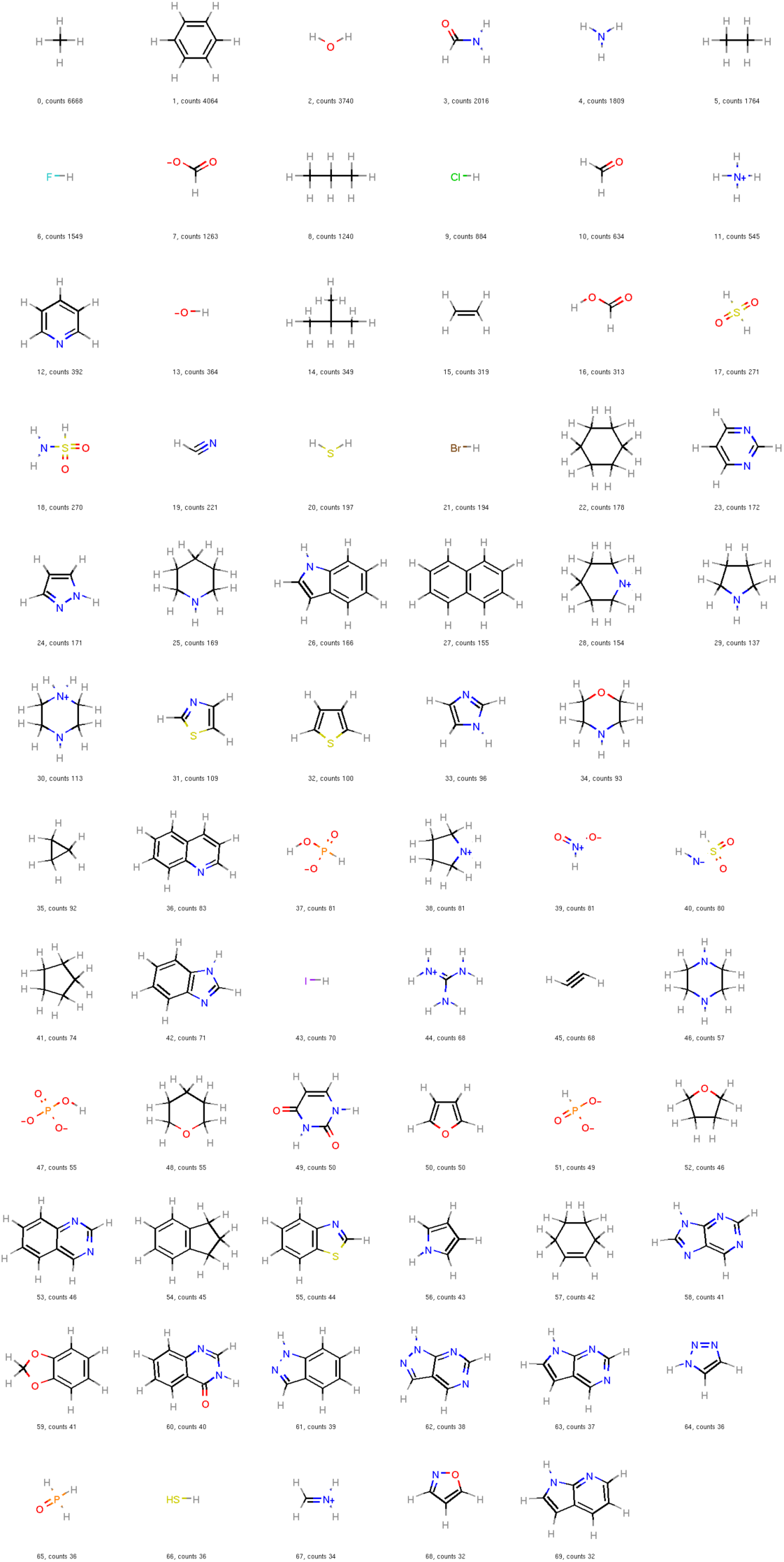
70 most frequently found fragments from the fragment library.

**Supplementary Figure 2.**
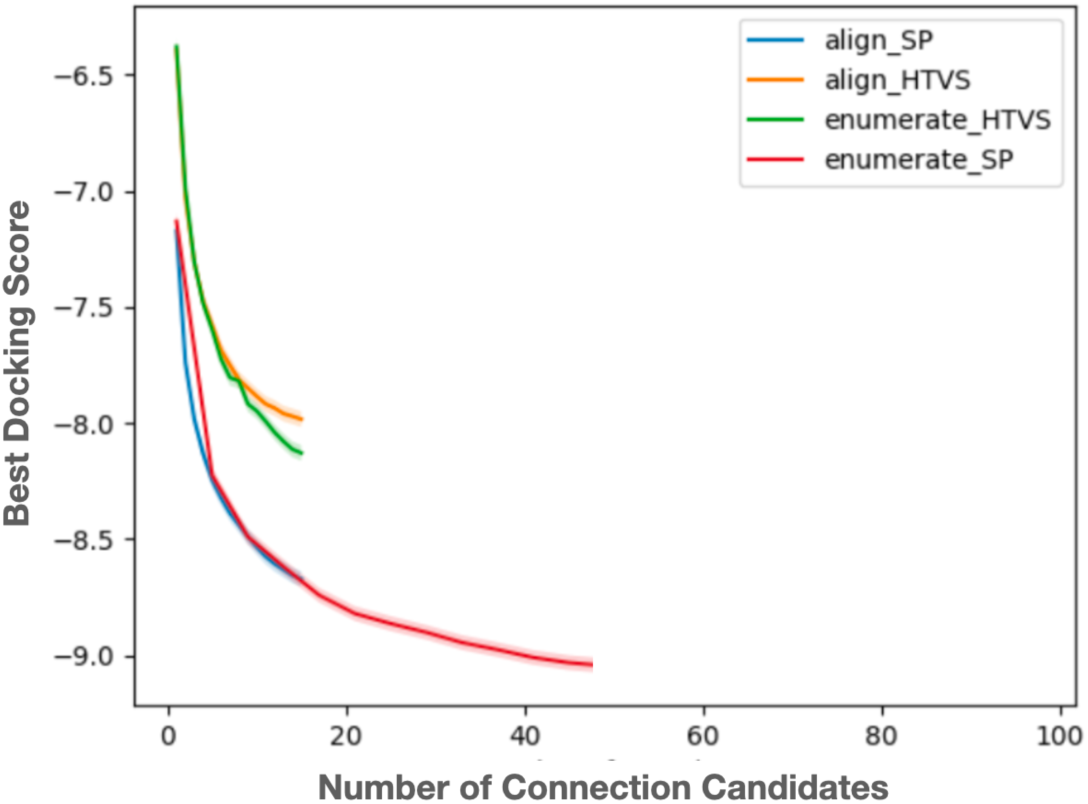
Docking score as a function of the number of connection candidates. For fixed sets of fragments, we sampled multiple chemically valid connections using our connection algorithm. Each resulting molecule was docked using either high-throughput virtual screening (HTVS) mode or standard precision (SP) mode. We report the best docking score as a function of the number of the sampled candidates. The results show that docking scores tend to plateau after sampling approximately 30–40 connection variants.

**Supplementary Figure 3:**
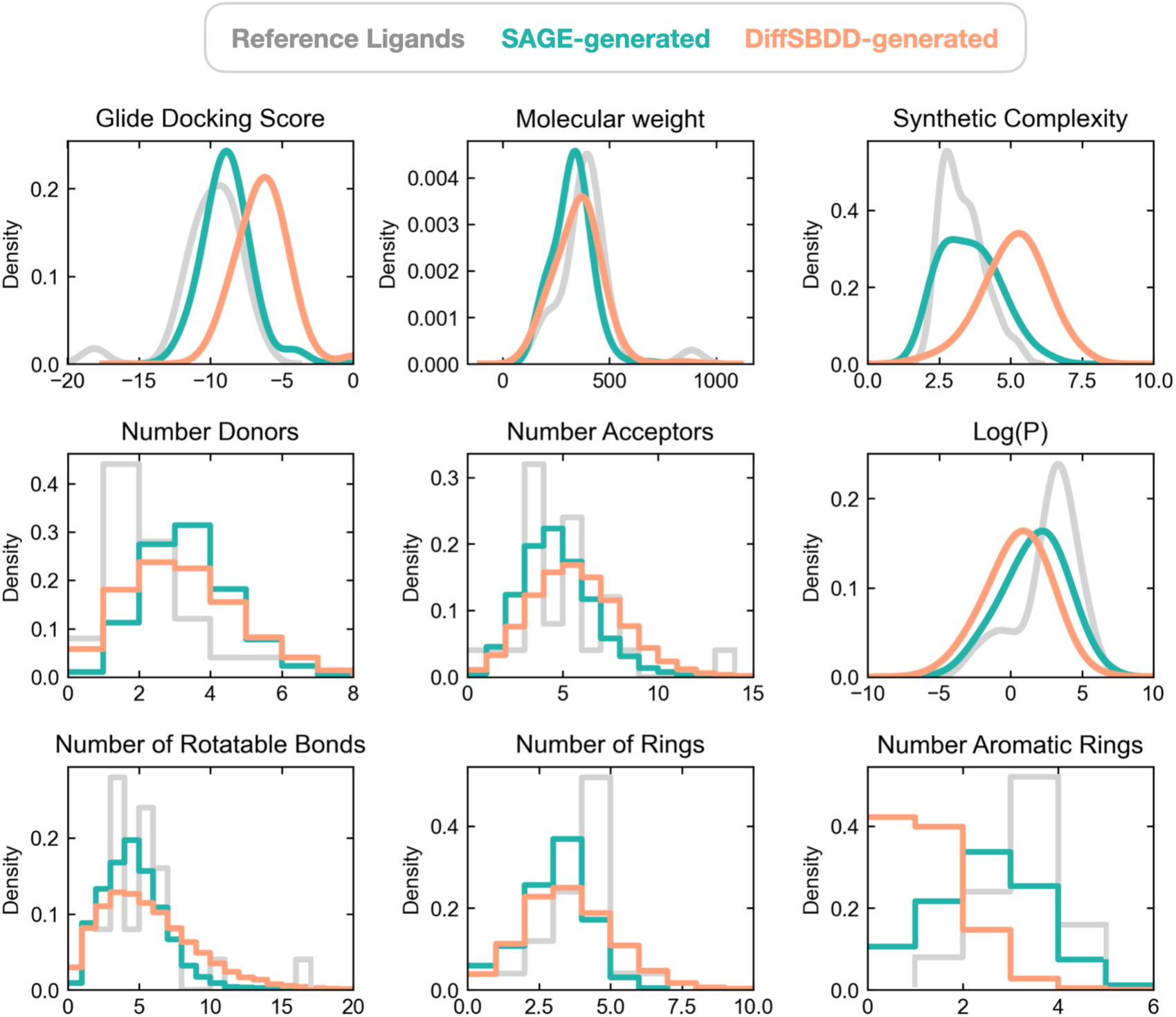
MedSAGE Produces Synthesizable and Drug-Like Molecules. For each target, several hundred ligands were generated using SAGE, using the 25 benchmarking target protein structures from the test set. The property distributions of the generated molecules (green) were compared to those of the known, high-affinity reference ligands (grey) and also ligands generated with all-atom diffusion method DiffSBDD (salmon). Distributions are plotted using kernel density estimations for continuous metrics and binned, normalized histograms for discrete metrics. Docking score are Glide scores at the target protein. Synthetic complexity score, calculated using the standard implementation in RDKit, is a relative measure of the ease of synthesizing the ligand, with lower scores indicating easier synthesis. All other properties are standard molecular descriptors computed with RDKit.

